# Cycles of gene expression and genome response during mammalian tissue regeneration

**DOI:** 10.1101/309989

**Authors:** Leonor Rib, Dominic Villeneuve, Viviane Praz, Nouria Hernandez, Nicolas Guex, Winship Herr, CycliX Consortium

## Abstract

**Background:** Compensatory liver hyperplasia — or regeneration — induced by two-thirds partial hepatectomy (PH) permits the study of synchronized activation of mammalian gene expression, particularly in relation to cell proliferation. Here, we measured genomic transcriptional responses and mRNA accumulation changes after PH and sham surgeries.

**Results:** During the first 10–20 hours, the PH- and sham-surgery responses were very similar, including parallel early activation of cell-division-cycle genes. After 20 hours, however, whereas post-PH livers continued with a robust and coordinate cell-division-cycle gene-expression response before returning to the resting state by one week, sham-surgery livers returned directly to a resting gene-expression state. Localization of RNA polymerase II (Pol II), and trimethylated histone H3 lysine 4 (H3K4me3) and 36 (H3K36me3) on genes dormant in the resting liver and activated during the PH response revealed a general *de novo* promoter Pol II recruitment and H3K4me3 increase during the early 10–20 hour phase followed by Pol II elongation and H3K36me3 accumulation in gene bodies during the later proliferation phase. H3K36me3, generally appearing at the first-internal exon, was preceded 5′ by H3K36me2; 3′ of the first-internal exon, in about half of genes H3K36me3 predominated and in the other half H3K36me2 and H3K36me3 co-existed. Further, we observed some unusual gene profiles with abundant Pol II but little evident H3K4me3 or H3K36me3 modification, indicating that these modifications are neither universal nor essential partners to Pol II transcription.

**Conclusions:** PH and sham surgical procedures on mice reveal striking early post-operatory gene expression similarities followed by synchronized mRNA accumulation and epigenetic histone mark changes specific to PH.

## Background

In developing multicellular organisms, cells proliferate and differentiate, and these processes are controlled by regulated gene expression. In embryonic development, many cells proliferate via a cell-division cycle under the control of cell-proliferation genes; subsequently, cells differentiate through the activation of differentiation-specific genes and cell-proliferation genes are generally silenced as cells often exit the cell-division cycle. The resulting quiescent differentiated cells possess unique sets of active and repressed genes. These sets of genes, however, can change as cells respond to physiological changes such as feeding or circadian cycles. Moreover, cells can re-enter the cell-division cycle as in the case of tissue regeneration.

In eukaryotes, both active and repressed genes are packaged in nucleosome-containing chromatin in which histones are reversibly modified — often reflecting the underlying gene transcription status. In this chromatin context, we study how cyclical programs of gene expression — i.e., circadian, nutrition and cell division — are regulated in differentiated cells using the mouse liver as model [1, 2, 3, 4]. We focus on gene expression as measured by RNA-transcript levels and study the relationship between gene occupancy by RNA polymerase II (Pol II) and two specific histone modifications: histone H3 lysine 4 trimethylation (H3K4me3) observed at active promoters and histone H3 lysine 36 trimethylation (H3K36me3) associated with the body of actively transcribed genes (reviewed in [5]). Here, we describe how these gene-expression markers change during liver regeneration.

By rapidly inducing proliferation of quiescent hepatocytes, the mammalian liver has a striking ability to compensate for cell loss caused by toxic substances or surgical removal [6, 7, 8]. Thus, for example, removal of 70% of the liver mass via partial hepatectomy (PH) leads to synchronous cell-division-cycle re-entry of most of the remaining hepatocytes. In mice, the first-round of hepatocyte division is accomplished within 60 h post PH; subsequent cell-division cycles together with cell growth lead to regeneration of the complete mass of the liver — compensatory hyperplasia — within two to three weeks [7, 8, 9]. We have used a characterized PH-induced mouse liver regeneration protocol [10] to study how a program of cell-division-cycle gene expression — dormant in the quiescent liver — is re-activated in the context of a differentiated tissue.

## Results

To analyze gene-expression changes associated with the cell-division cycle during liver regeneration, we integrated 70% PH into a dark/light and feeding-entrainment protocol described in [1] to study gene expression through the circadian cycle. Prior to 70% PH, mice were entrained for two weeks on 12 h dark/12 h light cycles with food provided only during the dark (waking) period (see Additional File 1: Figure S1a; and Additional File 10: Supplemental Material). PH was performed at Zeitgeber Time (ZT) 2, where ZT0 represents the beginning of the light/fasting period, and samples were collected at 1, 4, 10, 20, 28, 36, 44, 48, 60, and 72 h, as well as 1 and 4 weeks post PH (labeled X). To identify non-PH-related effects of the PH surgery, we performed parallel sham surgeries in which all procedures but the liver resection were included and collected samples at 1, 4, 10, 20 and 48 h post sham surgery (labeled S).

We measured gene expression (i) at the transcript level by ultra-high-throughput RNA-sequence determination (RNA-Seq) of poly(A)-selected RNA from individual livers and (ii) at the genomic level by measuring Pol II density using chromatin-immunoprecipitation on pooled sets of the three livers used for RNA-Seq analyses followed by ultra-high-throughput DNA-sequence determination (ChIP-Seq) as listed in Additional File 1: Figure S1b.

As illustrated in the hierarchical clustering dendrogram shown in Fig. 1a, the RNA-Seq analysis revealed that the triplicate livers had highly similar patterns of transcript abundance — in about three-quarters of cases the triplicates were immediate neighbors. Even the three separate sets of triplicate 0 h time points (C0), although clustering separately, displayed Pearson correlations of 0.94 or better (Additional File 1: Figure S1c). Globally, the clustering dendrogram revealed three groups of samples (labeled I, II, and III): Group I represents samples similar to the resting 0 h C0 time point, Group II includes 4, 10, 20, and 28 h post-PH samples along with 4 and 10 h sham samples, and Group III represents 36–72 h post-PH samples. Importantly, replicate samples never fell into different groups I–III, indicating that the PH protocol [10] was highly reproducible. Thus, in the ChIP-Seq analyses described below, where we pooled samples to have sufficient material for analysis, the signals were probably not significantly blurred.

**Figure 1.**
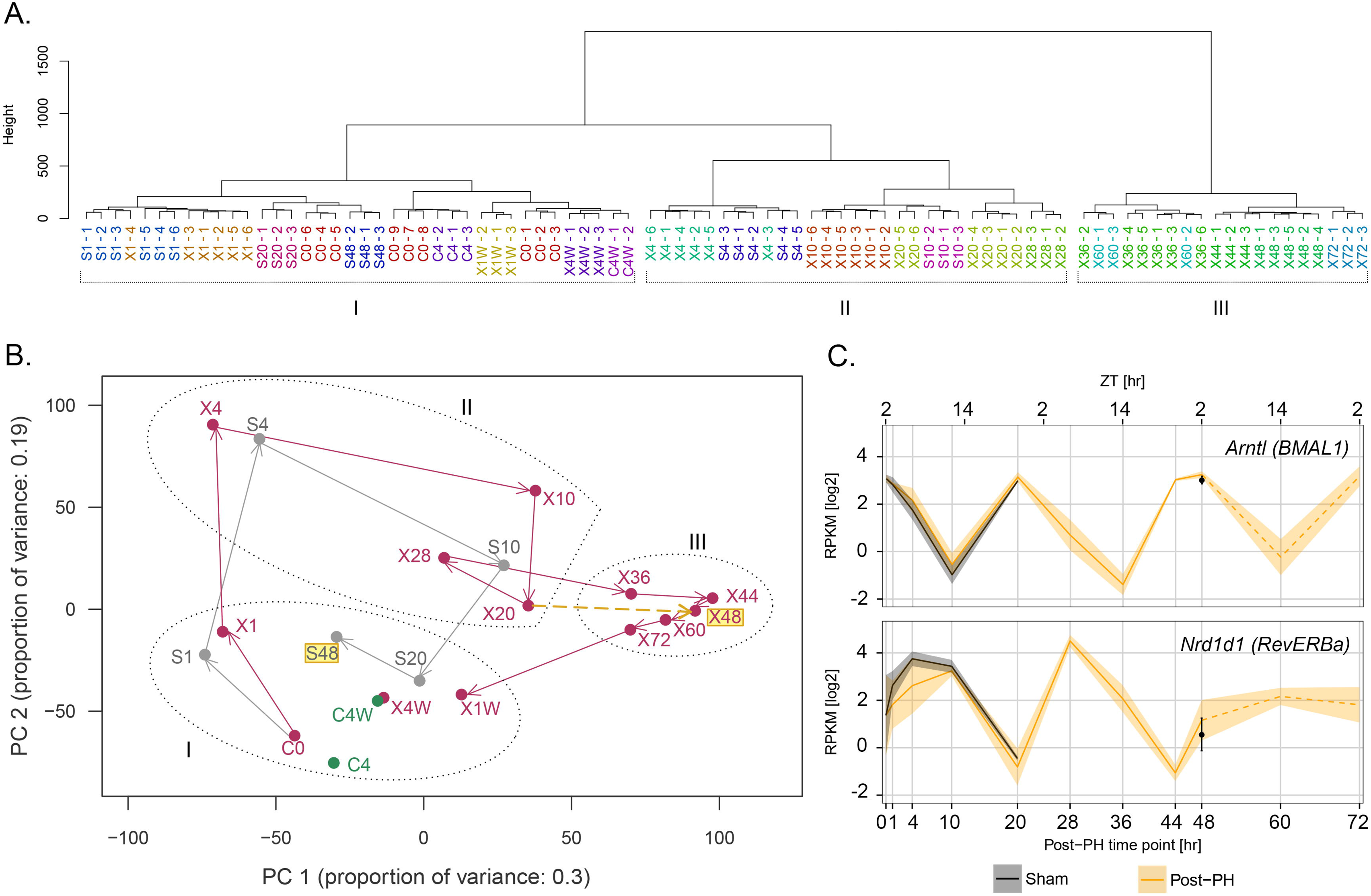
Gene expression profiles after sham and PH surgery. (**A**) Dendogram of hierarchical clustering of gene expression profiles for individual post-sham and post-PH samples. The 12,025 Set 1 and Set 2 genes listed in Additional Files 6 and 7: Tables S1 and S2 were used in the analysis. Replicate samples are color-coded. Sample numbers represent hours post treatment unless specified as weeks (W). C, control (no treatment); S, sham; X, PH. Branch lengths correspond to gene-expression differences among samples. The three principal branches are numbered I, II, and III, and set apart with brackets. (**B**) Two-dimensional PCA plot for components PC1 and PC2 of samples shown in part (A). Samples labeled C, S and X as in (A) are shown in green, gray, and red, respectively. The coordinates of replicate samples were averaged and are displayed as single dots; see Supplemental Figure S1D for standard deviations. Arrows indicate the paths followed by the post-sham (gray) and post-PH (red) samples. The S48 post-sham and X48 post-PH time points are highlighted in yellow and the X20 and X44 post-PH time points are connected by a dashed yellow arrow to emphasize the difference between the post-sham S20 to S48 and post-PH X20 to X48 trajectories in the PCA plot. Samples from branches I, II and III from the hierarchical clustering dendogram (Fig. 1A) are each indicated with dotted circles. (**C**) Expression profiles of the mouse core circadian-cycle genes *Arntl* and *Nrd1d1*; the corresponding human gene names *BMAL1* and *RevERBa* are given in parentheses. Log2 of RPKM quantifications for averaged replicate post-sham (black) and post-PH (yellow) samples are shown over 72 h; the shaded areas represent standard deviations. The hours post surgery are shown on the lower x-axis and ZT hours on the upper x-axis.

### The post-PH gene expression profile reveals two periodic cycles: an early cycle shared with sham-surgery mice followed by a PH-specific cycle

To identify stage-specific changes in gene expression, we used principal component analysis (PCA) to maximally differentiate patterns of transcript abundance amongst samples. The PCA revealed first (PC1) and second (PC2) principal components with 0.30 and 0.19 proportions of variances, respectively, thus accounting for nearly 50% of the variation in the samples. Fig. 1b shows a PCA plot for the aggregated results of each time point and treatment (see Additional File 1: Figure S1d for standard deviations). The analysis reveals a robust initial response over 4 h that is shared by both the PH and sham samples (compare X1 to S1, and X4 to S4), whereby both the PH and sham samples transit from Group I to Group II of Fig. 1a. The PH vs. sham similarity becomes progressively less pronounced at 10 h to 20 h (X10 vs. S10, and X20 vs. S20) and yet there is an evident shared clockwise trajectory — or “cycle” — for both the PH and sham samples. The 20- and 48-h sham samples return to the resting state Group I (Fig. 1a). In contrast, the PH samples form a new PH-specific “cycle” progressing from Group II (X20 and X28) to Group III (X36, X44, X48, X60, and X72) before returning to Group I, where they remain at one to four weeks post PH (X1W and X4W). The difference in change of trajectory between the sham and PH samples is exemplified by the comparison of the S20-to-S48 vector (right to left) to the X20-to-X48 vector (left to right). Thus, the PCA of the post-PH-gene expression profiles reveals two apparent cycles: an initial approximately 10–20-h cycle shared with mice subjected to sham surgery followed by a longer PH-specific cycle.

The similar 1–4 h sham and PH PCA responses were not a normal aspect of changing liver-gene expression owing, for example, to the circadian cycle, because the PCA position of samples from mice of the same ZT6 as the X4 and S4 mice but not manipulated (called C4) was very similar to the C0 sample (Fig. 1b). Thus, the sham and PH samples share an early gene-expression response that is dependent on the surgical manipulation but independent of the liver resection itself. This response might reflect the administered anesthesia and/or painkiller, skin wounding, or stress.

After four weeks, the post-PH (X4W) and untreated control (C4W) mice appear essentially identical in the PCA plot (Fig. 1b). Thus, there were no evident long-term effects of PH on liver gene expression.

### Unperturbed expression of two circadian cycle master regulators post PH

Although, as indicated by the PCA in Fig. 1b and the analyses described below, the PH procedure induces two robust cycles of gene expression, we found that important elements of a separate underlying gene expression cycle — the circadian cycle — remained unperturbed. Thus, consistent with previous observations [9] and as shown in Fig. 1c, the different cyclic transcript abundance profiles of the two circadian-cycle master regulators *Arntl* (*BMAL1* in human) and *Nrd1d1* (*RevERBα* in human) continued unabated after sham surgery or PH-induced liver regeneration. These PH results emphasize the robust nature of the circadian cycle of gene expression during liver regeneration.

### The post-PH-specific gene expression pattern displays a robust *de novo* cell proliferation component

To examine the post-PH gene expression changes in more detail, we divided the transcript levels corresponding to the 37,991 individual Ensembl 67/NCBI37 protein-encoding and non-coding genes into three sets representing (i) transcripts that were not detected in any sample (Set 1); (ii) transcripts that did not vary significantly during the course of PH recovery (Set 2); and transcripts that clearly varied (Set 3) as indicated in Table 1. The undetected Set 1 transcripts were either owing (i) to their absence in the liver and/or (ii) to the experimental selection for poly(A) transcripts (e.g., the non-polyadenylated replication-dependent histone-encoding transcripts went undetected). Consistent with a loss of non-poly(A) transcripts, 32% of Set 1 transcripts were non-protein encoding as opposed to less than 10% of Sets 2 and 3 transcripts.

**Table 1.** Transcript detection and characterization.

| **Set** | **N° genes** | **% of genes in set**<br>**protein coding** | **% of KEGG 124**<br>**cell cycle genes** |
| --- | --- | --- | --- |
| 1. Gene transcripts not detected | 25966 | 68% | 9% |
| 2. Gene transcripts detected and stable | 6528 | 93% | 32% |
| 3. Gene transcripts detected and changing | 5497 | 95% | 59% |

As Set 3 transcript levels vary, we could identify subsets of co-varying gene-transcript profiles using the Partitioning Around Medoids (PAM) clustering algorithm. The algorithm chooses one representative gene, called ‘medoid’, for each of any defined number of groups, and iteratively adds genes to each group minimizing the dissimilarities among genes within each group. To minimize differences within groups, as the iterative process proceeds, medoids can be replaced by more representative genes if the exchange decreases the overall dissimilarity among genes within a group or cluster. Here, we used the algorithm to probe transcript-abundance-profile similarity.

Overall, gene-transcript levels were seen to either decrease or increase during the course of PH recovery before returning to the starting levels. After an initial PAM analysis defining from 2 to 30 subsets (see Additional File 2: Figure S2a for the 2–13 subset analyses), we selected for further analysis a seven-subset grouping (called Set 3.1 to Set 3.7) where the size varied from 444 (Set 3.3) to 1060 (Set 3.5) genes (Fig. 2a). In subsets Set 3.1 to Set 3.3 gene-transcript levels decreased overall (bracket I) and for Set 3.4 to Set 3.7 transcript levels increased overall (bracket II) following PH.

**Figure 2.**
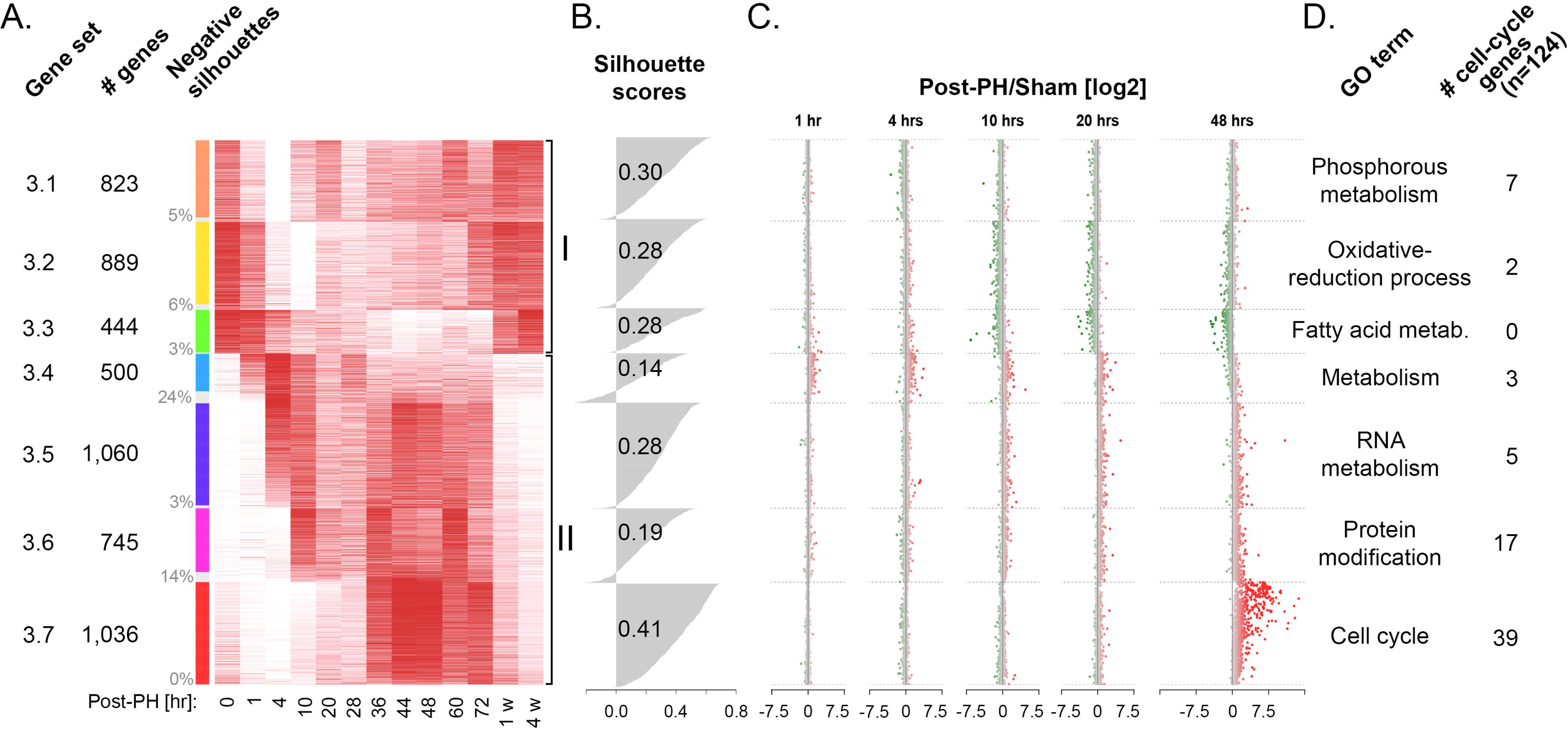
Changing post-PH gene-expression patterns and comparison with sham-surgery samples. (**A**) Heat-map display of seven-set PAM clustering results for transcripts that varied post-PH gene-expression patterns (i.e., Set 3). The individually normalized relative post-PH transcript abundance (red, high; white, low) for each gene is shown. The seven PAM-clustering sets (Set 3.1 to Set 3.7 indicated by the color coding column to the left) are each grouped together with the comparative medoid at the top of each set and decreasing gene-expression similarity shown from top to bottom. Set 3 PAM-clustering subset name (column 1), number of genes per subset (column 2), and percentage genes per subset with a negative silhouette clustering score (column 3) are given. PAM-clustering sets with genes down-regulated (Set 3.1 to Set 3.3) and up-regulated (Set 3.4 to Set 3.7) post PH are indicated by the brackets labeled I and II, respectively. (**B**) Silhouette-score distributions and averages for each PAM-clustering subset. (**C**) Transcript-abundance comparison between post-PH and post-sham samples at 1, 4, 10, 20 and 48 h. Genes are indicated as dots in the same order as in (A). The post-PH/sham ratio is given in log2 scale (x axes). Positive (higher post-PH expression in red) and negative (higher post-sham expression in green) log2 scores are indicated as color gradients. Gene transcripts with post-PH and post-sham log2 RPKM quantifications less than 0 are not shown. (**D**) Predominant specific function of genes for each Set 3 PAM-clustering subset and number of genes per cluster included in the 124-gene *Mus musculus* KEGG cell cycle pathway. Only the most representative GO term that is specific for an individual subset is listed. The full list of enriched GO terms is given in Additional File 8: Table S3.

Fig. 2a shows a heat map of the Set 3.1 to Set 3.7 subsets. The gene-transcript levels measured as <u>r</u>eads <u>p</u>er <u>K</u>b of transcript per <u>m</u>illion mapped reads (RPKM) were individually normalized to their maximal (red) and minimal (white) levels. For each subset, the individual and average gene-transcript “silhouette scores” (i.e., the degree of similarity to the subset-specific medoid vs. dissimilarity to the six other subset-specific medoids) are given in Fig. 2b. Subsets can have gene transcripts with negative silhouette scores. Thus, Set 3.4 (24%) and Set 3.6 (14%) subsets display prominent negative silhouette score groups, whereas Set 3.7 has only one negative silhouette transcript out of 1036 gene transcripts. Indeed, Set 3.7 is special in that it has a significantly higher average silhouette score (0.41) than the other Set 3 subsets, and a major Set 3.7-like subset was evident across the 2–13 group PAM selection (Additional File 2: Figure S2a). It was also most active at 36 to 72 h post PH and is enriched in cell proliferation genes (see below).

Set 3.7 is also special in that it is PH specific, as shown by the sham and post-PH sample comparison in Fig. 2c. Here, with the same Fig. 2a gene order, post-PH-to-sham transcript level ratios at 1, 4, 10, 20 and 48 h are shown. Consistent with the early, shared PCA-plot PH/sham cycle (Fig. 1a), the 1 and 4-h time points display little PH-to-sham difference. But by 10–20 h, particularly in the Set 3.3 subset, there is an evident over representation of some gene transcripts in the sham samples (shown in green); this Set 3.3 effect becomes even more prominent at 48 h. By 20 h, the sham sample has returned to the group I baseline (see Fig. 1b), ending the shared PH/sham PCA cycle. These transcripts may represent genes involved in liver functions that remain overall lower in the regenerating liver as some cells become dedicated to cell proliferation. In contrast, the genes in the Set 3.4 to Set 3.7 subsets have generally higher corresponding transcript levels in the PH than sham samples (shown in red). This PH-specific enhancement is particularly prominent with Set 3.7, consistent with its high activity at 36 to 72 h post PH (Fig. 2a).

### The PH-specific Set 3.7 is rich in cell-cycle-related genes

To probe the overall biological-function enrichments in the different gene sets, we probed the Gene Ontology database (GO; [11]). We thus statistically annotated the Set 1, Set 2, and Set 3 gene sets, as well as Set 3.1 to Set 3.7 subsets (GO classifications, or terms, with an adjusted p-value (p) of less than p = 0.05 as listed in Additional File 6: Table S1). Additional File 2: Figure S2b shows REVIGO [12] graphic representations of Set 1 through Set 3, in which the semantic relationship of GO terms with p < 10^−10^ are displayed with a p-value color scale. Thus, each cluster of terms (i.e., cluster of dots) is generally functionally related. To illustrate the functional enrichments, representatives with low p-values from different semantic clusters are annotated, with their p-values given. As expected, the non-expressed Set 1 gene set is devoid of terms specifically involved in liver function. In contrast, both Set 2 and Set 3 are enriched in terms involved in liver-specific functions as well as general cellular and metabolic functions, with Set 3 also possessing many terms associated with cell proliferation (e.g., cell cycle p = 3.9 × 10^−77^; cell division p = 3.8 × 10^−40^).

To probe Set 3 further, we performed an individual gene enrichment analysis for each of its Set 3.1 to 3.7 “subsets.” For Set 3.1 to Set 3.6, the most statistically significant terms were generally related to cell metabolism, growth, and regulation as indicated in Fig. 2d; Set 3.7 stood out by its large number of highly significant cell-cycle-related terms as shown in Table 2. For three of the 13 terms listed, the term was not found to be more significant than p = 0.01 in any other Set 3 subset; for ten of the 13 terms shown, the term scored second best in Set 3.6 with the p-values listed in Table 2. The cell-cycle relationship of Set 3.7 is strong and is consistent with its peak expression between 36 to 72 h, when mouse hepatocytes are rapidly proliferating [9].

**Table 2.**
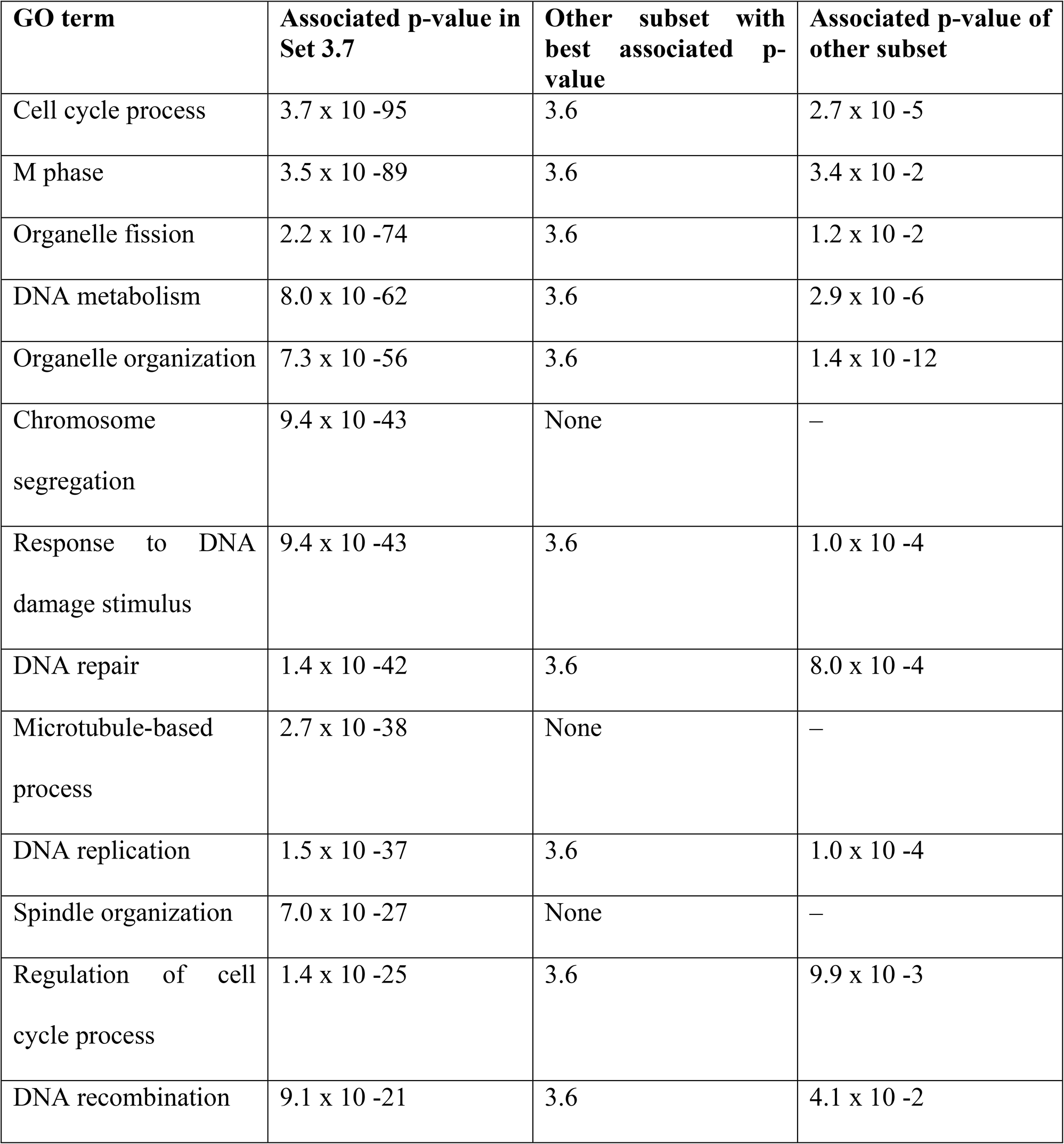
Predominant Set 3.7 GO terms with their associated p-values compared to the best associated p-value for the same specified GO term in Set 1 – Set 3.6.

### Similarity and divergence in cell-division-cycle-gene response to sham and PH surgeries

To probe the cell-division-cycle relationship of different gene sets further, we studied the expression patterns of the 124 genes listed in the KEGG cell-division-cycle pathway. We first mapped them to the different sets: 11 (9%) mapped to Set 1, 40 (32%) mapped to Set 2, and 73 (59%) mapped to Set 3 (Table 1). Of the 73 Set 3 cell-division-cycle genes, 39 (over 30% of all KEGG cell-cycle pathway genes) mapped to Set 3.7 and 17 to Set 3.6, with single digit numbers in the five other Set 3 subsets (Fig. 2d). Thus, Set 3.7 followed by Set 3.6 are the most highly cell-division-cycle-related co-varying gene expression sets.

We then examined directly the activities of the KEGG cell-cycle pathway genes in the sham and PH samples. Fig. 3 shows a heat map of absolute RNA levels over the post-sham and -PH time courses in which the genes are ordered according to their RNA levels at time 0 h. Whereas some specific RNAs display constant levels over the course of PH-induced liver regeneration (e.g., *Skp1a, Rb1, E2f5*), the majority varied, and usually over the course of 36 to 72 h (e.g., *Ccna2, Ccnb2, Ccne2* cyclin genes, *Cdc20, Plk1, Bub1, Cdc25c* M-phase genes). Of note, RNAs corresponding to the *Ccnd1* gene (see arrow), encoding the G1-phase regulator Cyclin D1, peak at three separate time points, 10, 36, and 60 h. Strikingly, even for cell-division-cycle genes, the pattern of RNA-level variation in the sham samples — where few cells proliferated [10] —parallels that of the post-PH samples up to 20 h, with Spearman correlations between 0.99 and 0.96; at 48 h, however, the high similarity is lost (0.64 Spearman correlation).

**Figure 3.**
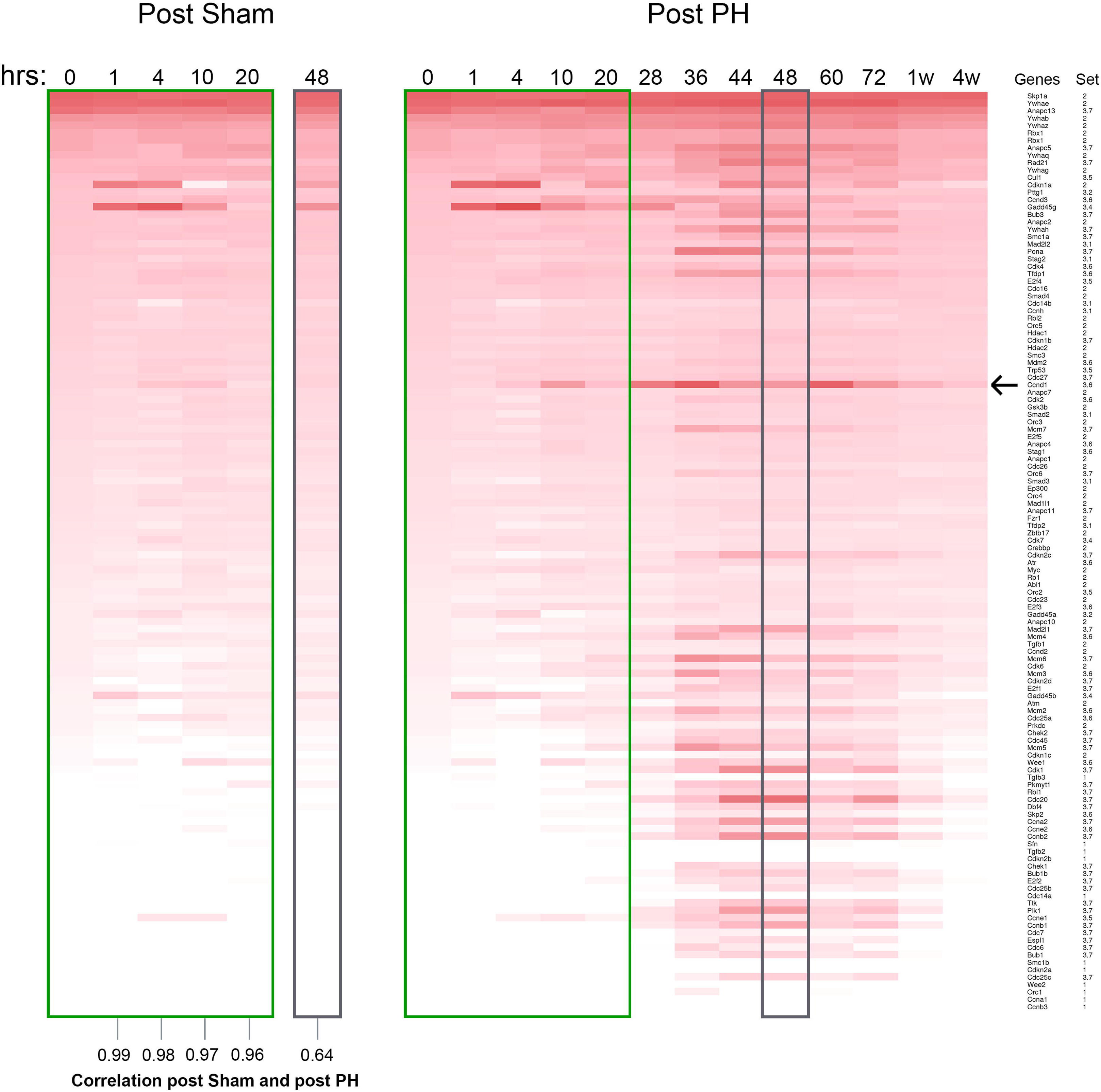
Transcript abundance changes of the 124 *Mus musculus* KEGG cell-cycle genes post-sham and post-PH. The 124 *Mus musculus* KEGG cell-cycle genes are organized in a heat map according to their 0 h log2 RPKM transcript level (high to low, top to bottom). The 0 h log2 RPKM transcript abundance level is compared separately to the post-sham (left) and post-PH (right), using gene-specific z-scores. Gene names and associated Set 3 PAM-clustering subset are listed to the right. Shared post sham and PH 0 to 20 h and 48 h samples are outlined in green and gray respectively. Arrow, *Ccnd1* gene.

In Additional File 2: Figure S2c and more dramatically in Additional File 9: Movie S1, we have superimposed the RNA levels over time of the KEGG cell-cycle pathway genes on the KEGG cell-cycle pathway rendered by Pathview [13]. Where the KEGG cell-cycle Pathway view lists more than one gene per node (with the six-membered ORC and MCM complexes shown in a separate box), one representative gene (see Additional File 2: Figure S2d) was selected for display in Additional File 2: Figure S2c and the movie. The displays shown in Additional File 2: Figure S2c and Additional File 9: Movie S1 illustrate the coordinated pattern of cell-division-cycle gene expression during liver regeneration.

### Genomic responses to PH

Having documented gene expression during liver regeneration post PH by RNA-Seq of poly(A)-containing RNAs, we turned to the genomic response to PH by ChIP-Seq analysis. We paid particular attention to the transcriptional reactivation of cell-proliferation genes that have been largely quiescent in the resting adult liver prior to PH. As in the RNA-Seq analyses, the ChIP-Seq results of the sham samples paralleled those of PH and thus are not specifically described below.

We studied the relationships of Pol II occupancy at the promoter (defined as 250 bp upstream and downstream of the annotated transcriptional start site, TSS) and in the “body” of the gene transcription unit (defined as 500 bp downstream of the TSS to 2 kb downstream of the poly(A)-addition signal) with H3K4me3 at the promoter (500 bp upstream and downstream of the TSS) and H3K36me3 analyzed as for Pol II in the gene body. Fig. 4 shows such a comparison with two genes: the *Cxxc1* and *Ska1* genes over a 40 kb region. These genes were selected for display because (i) they are divergently transcribed but clearly non-overlapping and (ii) one, *Cxxc1*, encodes a CpG-binding subunit of the Set1 H3K4 methyltransferase whose corresponding mRNA is found in the non-varying RNA-Seq Set 2 and the other, *Ska1*, encodes a subunit of a microtubule-binding subcomplex of the outer kinetochore involved in chromosome segregation during mitosis and whose corresponding mRNA is found in the RNA-Seq Set 3.7.

**Figure 4.**
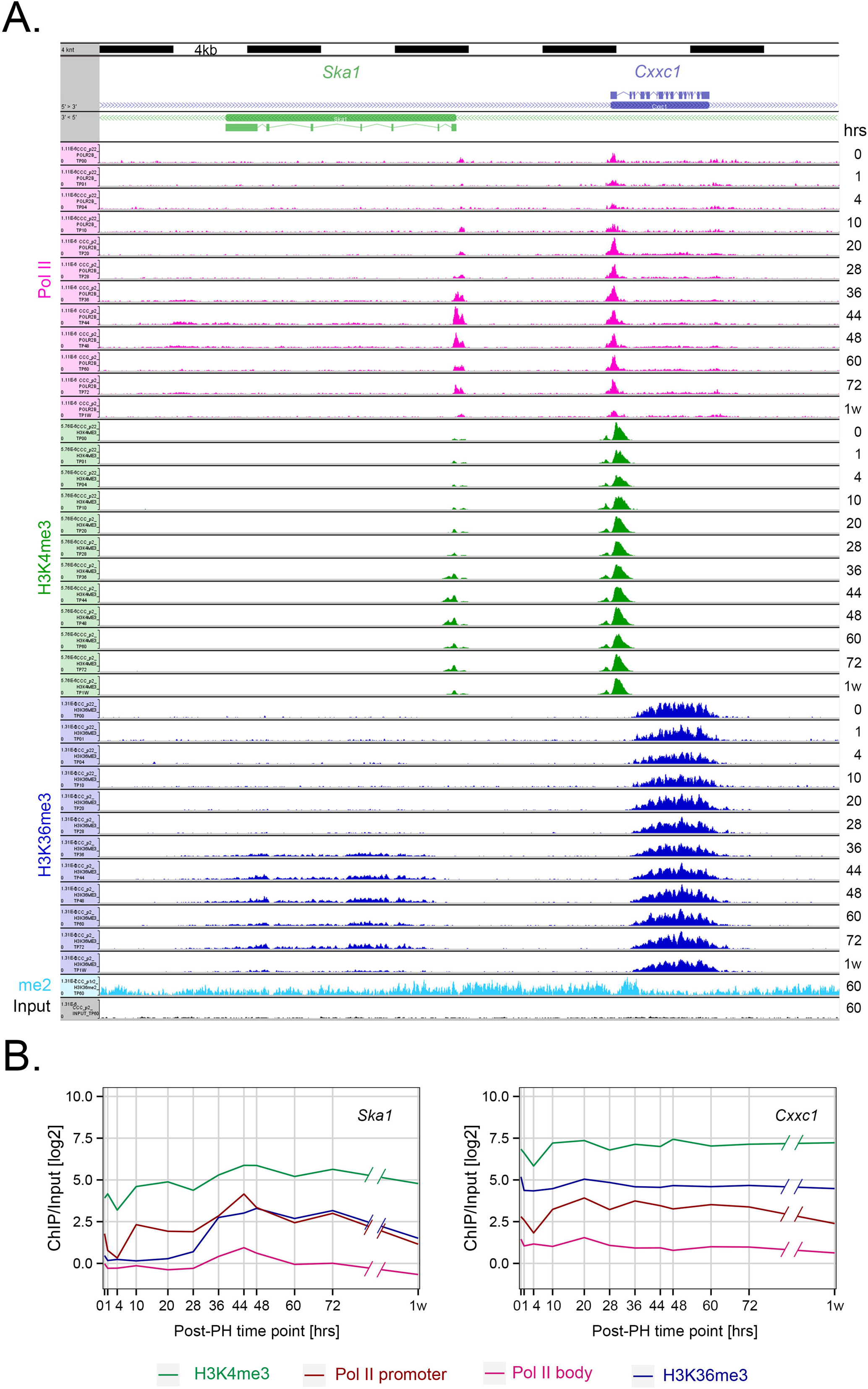
Post-PH genomic responses for the divergently transcribed PH-induced *Ska1* and non-PH-induced *Cxxc1* genes. (**A**) Genomic view of the *Ska1* (left) and *Cxxc1* (right) genes. Densities of the central 50 bp of paired-end reads for Pol II (pink), H3K4me3 (green) and H3K36me3 (blue) ChIP fragments are shown for 0 h to 1 week post PH. Similarly, densities for H3K36me2 and input fragments at 60 h post PH are shown. (**B**) H3K4me3 (green), Pol II promoter (burgundy) and body (pink), and H3K36me3 (blue) log2 ChIP/Input fragment-density comparison for the *Ska1* (left) and *Cxxc1* (right) genes post PH. The central 50 bp of the paired-end reads were used for quantification. The regions used for each quantitation are given in the text.

Consistent with the RNA-Seq results, the levels of Pol II in the transcription unit body were relatively stable for the *Cxxc1* gene but increased during 36 to 48 h for the *Ska1* gene (Fig. 4b). Although there is some decrease in all four signals at 4 h in the *Cxxc1* gene, particularly for promoter Pol II, H3K4me3 and H3K36me3, the levels are relatively stable from 10 h on. In contrast, for the *Ska1* gene the four signals vary significantly during the course of the post-PH process, where H3K4me3, Pol II at the promoter and in the gene body exhibit a wave of presence at 36 to 48h, with its maximum at 44 and 48h. H3K36me3 initially follows this pattern but stays high longer, until 72h. Additional File 3: Figures S3a–d show other gene-pair examples representing Set 2 and Set 3 mRNA transcript profiles; Additional File 3: Figure S3e shows the cell-proliferation gene *Mki67*. They reveal patterns similar to those in Fig. 4.

To study more generally the patterns of Pol II occupancy and histone methylation on genes whose expression is activated *de novo* post PH (referred to as “Post-PH” genes), we used the robust RNA-Seq data sets to identify genes with low corresponding transcript levels in the ZT2 resting liver (i.e., 0 h) but with elevated levels at some point post PH. Because the transcript-level patterns and GO enrichment profiles for Set 3.4 to 3.7 vary, we analyzed the Post-PH genes of each set separately as shown in Fig. 5, and focus our description on those in Set 3.7 where Post-PH genes represent 30% (307 out of 1036) of all the genes in this cell-cycle-enriched gene set.

**Figure 5.**
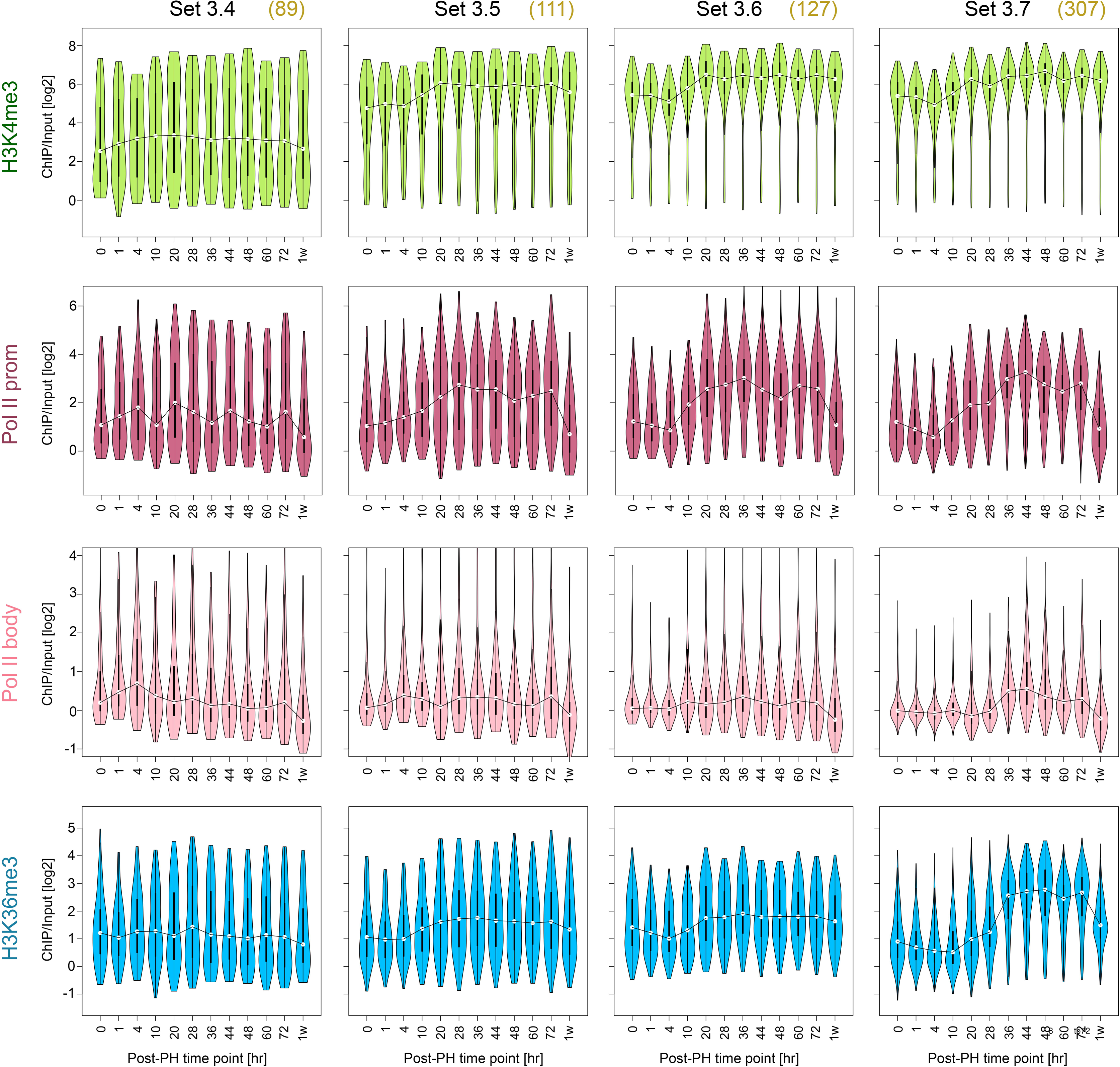
Post-PH genomic responses of genes activated by PH (Post-PH genes). H3K4me3 (green), Pol II promoter (burgundy) and body (pink), and H3K36me3 (blue) log2 ChIP/Input fragment-density ratios were determined as described in Fig. 4 legend and text. The numbers in parentheses represent the number of Post-PH genes in each of Sets 3.4 to 3.7 and thus those used in the analysis. In each “violin” plot the width (x-axis) of the display represents fragment density at each respective ChIP/Input density (y-axis). For each display, the line links the median for each distribution.

Fig. 5 shows the distributions of Post-PH genes for the four marks measured in Fig. 4 in the form of “violin” plots. We used the medians (linked in the figure) to make the following conclusions. The strongest signal was H3K4me3 followed by Pol II within the promoter, and then H3K36me3 and Pol II within the gene body. Furthermore, for Set 3.6 and particularly Set 3.7, the Pol II-promoter signal displays an initial dip at 4 h, followed by a two-step augmentation centered on 4 to 20 h followed by 28 to 44 h. Although less pronounced, a similar early increase in signal can also be discerned in the H3K4me3 signal. In contrast, in Set 3.7, both the Pol II-body and H3K36me3 signals display a predominant one-step augmentation at 28 to 44 h. Lastly, the two Pol II signals and, to a somewhat lesser extent, the H3K36me3 signal return to the starting level at one week. In contrast, the H3K4me3 signal remains high. These conglomerated patterns are all reflected in the *Ska1* Set 3.7 gene shown in Fig. 4.

Of separate note, the unusual bimodal pattern seen for each H3K4me3 time point in Set 3.4 is largely owing to many genes in this set being expressed at low levels. Indeed, some of these transcripts may represent RNAs from minor populations of non-hepatocytes in the regenerating liver. Consistent with this suggestion, Set 3.4 is the most enriched for the GO term “inflammatory response” (Additional File 8: Table S3).

In summary, Set 3.7 genes activated during PH-induced liver regeneration display the most coherent pattern for the four marks studied here, perhaps owing, at least in part, to their highly coherent gene expression profile (e.g., 0.41 silhouette score in Fig. 2b). This coherence, with the time points that we have analyzed, has allowed us to discern that for Set 3.7 Post-PH genes (i) Pol II within the promoter and to a lesser extent H3K4me3 display an overall coordinate bimodal 4–20 and 28–44 h activation pattern and (ii) Pol II within the gene body and H3K36me3 display a unimodal activation pattern coincident with the second of the Pol II promoter/H3K4me3 increases.

### Variable H3K36me2 to H3K36me3 transitions at the first internal exon of transcribed genes

Across the transcription unit, the H3K36me3 mark accumulates preferentially toward the 3′ end. Although H3K36me3 accumulation occurs on intronless genes [14] as shown for the *Cebpa* gene in Supplemental Fig. 4, H3K36me3 deposition is influenced by pre-mRNA splicing of intron-containing genes [14, 15]. Indeed, H3K36me3 deposition first appears around the first internal exon [14, 16]. We also observed such first-internal-exon H3K36me3 deposition patterns in our data sets. For example, Fig. 6a shows a set of three neighboring illustrative genes: *Txndc9, Eif5b*, and *Rev1* at 60 h post PH, with double-headed arrows indicating the position of the first internal exon for each gene.

**Figure 6.**
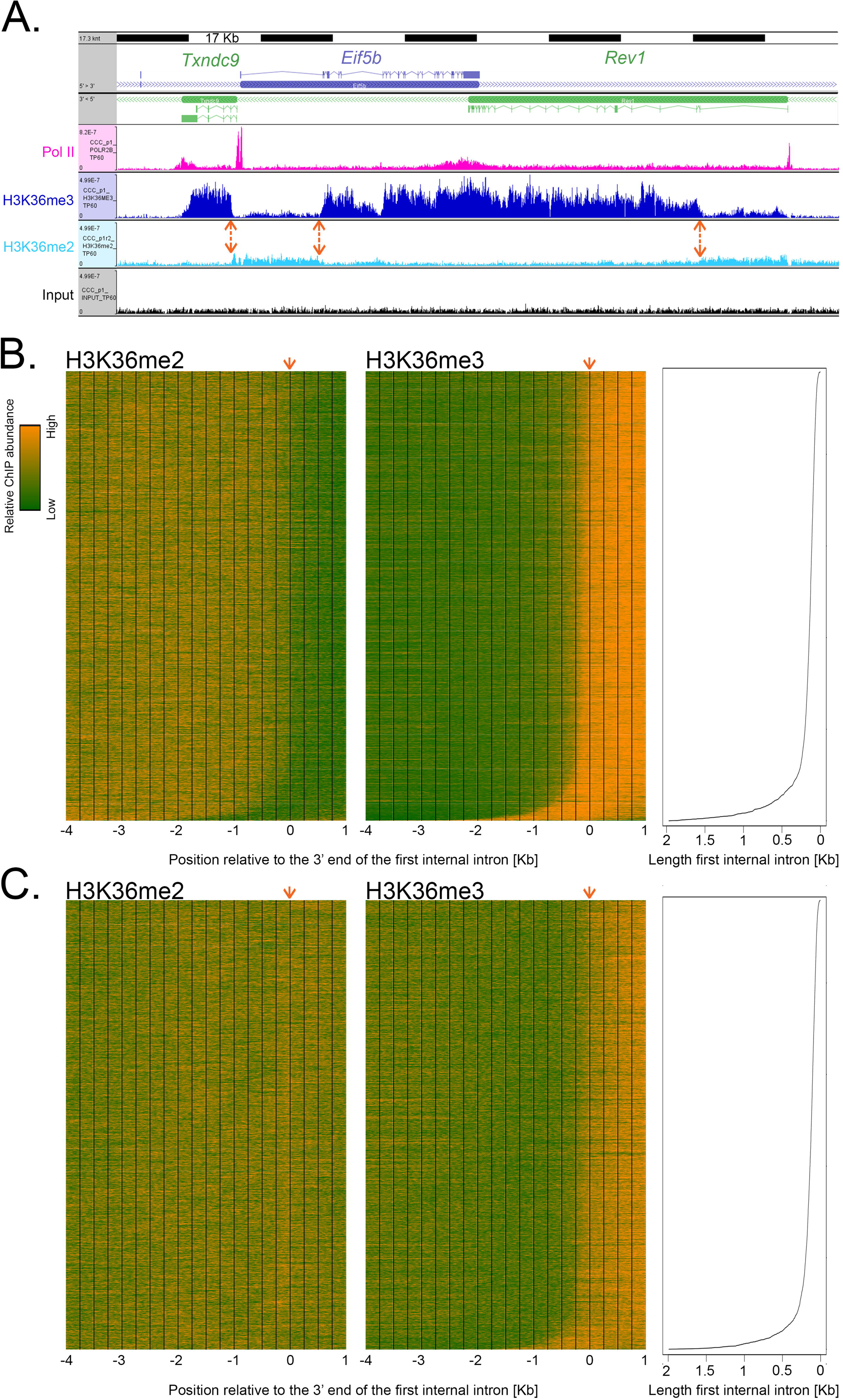
Relative accumulation of H3K36me2 and H3K36me3 marks in genes. (**A**) 170 Kb genome view of the *Txndc9* (antisense), *Eif5b* (sense) and *Rev1* (antisense) genes. Densities of Pol II (pink), H3K36me3 (dark blue), H3K36me2 (light blue) and Input (black) fragments are shown for samples at 60 h post PH as described in Fig. 4 legend. Double-headed orange arrows indicate the position of the 5′ end of the first internal intron of each gene. (**B**) and (**C**) Base-pair-resolution density of the central 50 bp of H3K36me2 (left) and H3K36me3 (right) ChIP fragments from 4 Kb upstream to 1 Kb downstream of the 3′ end of the first internal exon at 60 h post PH. Transcription units with a minimum of three exons (9801 total) were selected for analysis: the 4900 transcription units with higher first-internal exon H3K36me3 density at 60 h (see text) are shown in (B) and the remaining 4901 transcription units in (C). The orange arrows indicate the position of the 3′ end of the first internal exon used for alignment. The transcription units are sorted from top-to-bottom according to increasing first-internal-exon length, as indicated in the right-hand panels. Color scale: orange, high H3K36me2 or H3K36me3 density; green, low H3K36me2 or H3K36me3 density.

To determine whether this first-internal-exon-related H3K36me3 deposition was a general property of transcribed genes in the liver during regeneration, we prepared H3K36me3-density heat maps using the 60 h time point as shown in Fig. 6b and c. For this analysis, the H3K36me3 signal was measured over a 2 kb region extending downstream of the 5′ end of the first internal exon for each of the 9801 Set 2 and Set 3 genes (i.e., transcribed) with an internal exon (i.e., three or more exons); 8 genes with first-internal exons greater than 2 kb in length were removed from the analysis. The genes were separated into two groups, above (Fig. 6b) and below (Fig. 6c) the median, according to the level of H3K36me3 signal thus determined. In each case, the genes were aligned according to the position of the 3′ end of the first internal exon (position 0) and ordered from top to bottom according to the increasing length of the first internal exon. This organization had the effect of moving the 5′ end of the first-internal exon to the left from top to bottom as indicated in the adjoining line diagram.

The H3K36me3 heat maps make two clear points. First, the pattern of H3K36me3 signal (shown in orange) is similar for both the upper and lower quartile sets, although the signal is naturally more robust for the set above the median, and, second, the transition to H3K36me3 signal aligns with the shifting position of the 5′ end of the first internal exon. These results are consistent with a significant global signal for H3K36me3 deposition at the first-internal exon of transcribed genes [14, 16].

H3K36me3 deposition requires H3K36me2 modified histones. In metazoans, H3K36me2 modification is associated with actively transcribed genes and is performed by multiple enzymes, whereas only one methyltransferase, SetD2, is responsible for H3K36 di-to-trimethylation (reviewed in [17]). To determine the cross-talk between these two closely-related H3K36 modifications, we probed the spatial relationship between genomic H3K36me2 and H3K36me3 modifications. Profiles for this ChIP-Seq are shown in Fig. 4a, 6a, and 7a, and Additional File 3: Figure S3. We observed an elevated H3K36me2 signal at transcribed genes. Fig. 6b and c (left) show heat maps for the H3K36me2 signal as done for H3K36Me3. Surprisingly, the H3K36me2 signal pattern differed for the two sets above (Fig. 6b) and below (Fig. 6c) the median. For the upper half, there was a very evident decrease in H3K36me2 signal at the point at which the H3K36me3 signal appears at the first internal exon. In contrast, for the lower half, the levels of H3K36me2 signal remain essentially constant as the H3K36me3 signal appears at the first internal exon. As the lower and upper H3K36me3 densities were defined over the first internal exon region, the difference in H3K36me2 pattern may reflect a more pronounced, perhaps even complete, trimethylation of H3K36 in the upper gene set. In any case, H3K36me2 appears to establish a platform at transcribed genes for H3K36me3 methylation starting at the first intron-first internal exon junction and extending toward the end of the gene.

**Figure 7.**
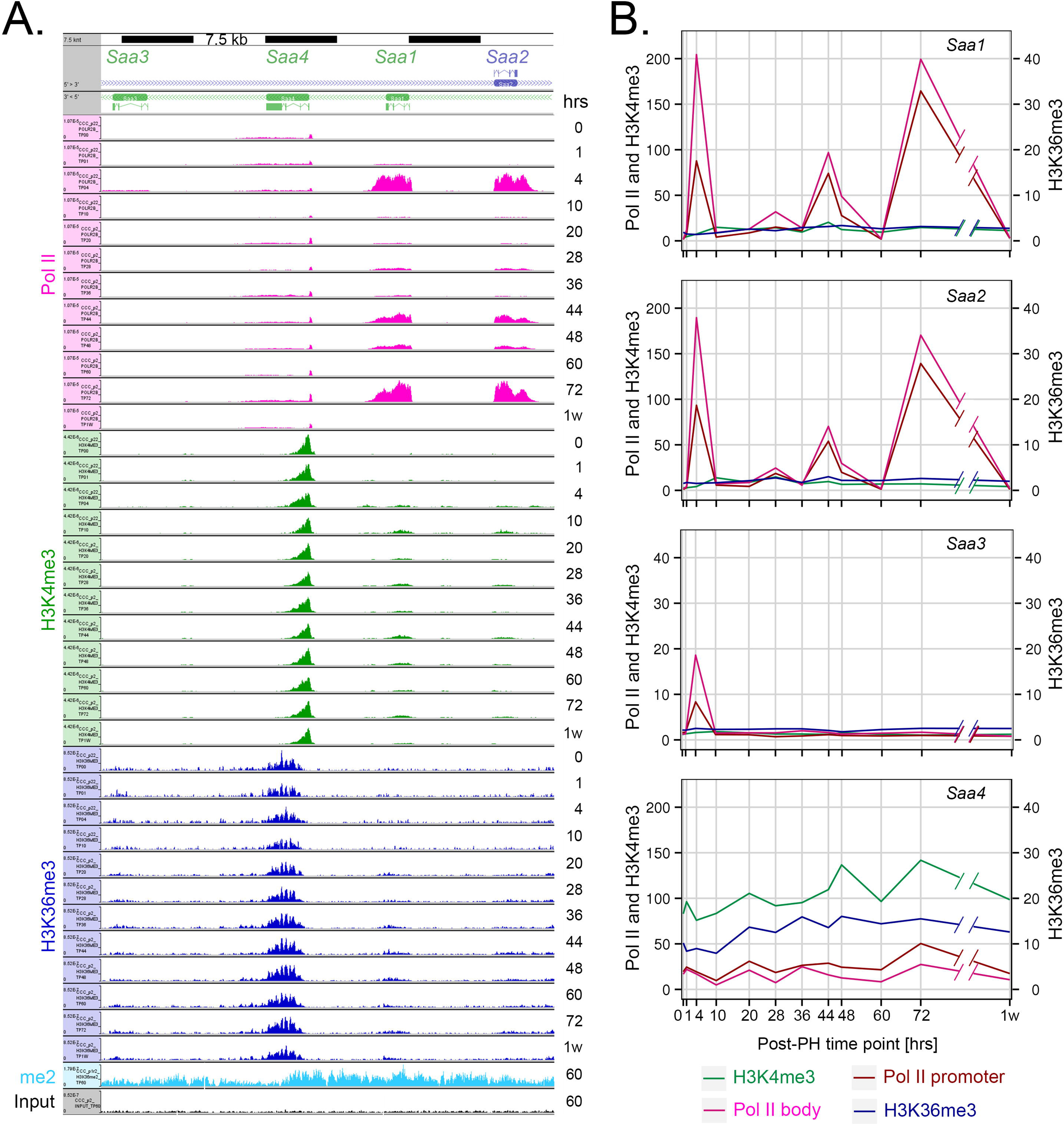
Genomic responses of the acute-response *Saa* genes post PH. (**A**) 45 Kb genome view of the *Saa1* to *Saa4* genes. Visualization of Pol II (pink), H3K4me3 (green), H3K36me3 (dark blue), H3K36me2 (light blue) and input (black) fragment densities is as described in Fig. 4 legend. (**B**) H3K4me3 (green), Pol II promoter (burgundy) and body (pink), and H3K36me3 (blue) log2 ChIP/Input fragment-density comparison for the *Saa1, Saa2, Saa3*, and *Saa4* genes (top-to-bottom) post PH. The genomic regions and ChIP-fragment sequences (central 50 bp) used for each quantitation are as in Fig. 4. The y-axis scales for levels of Pol II and H3K4me3 (left), and H3K36me3 (right) differ.

### Genes can display high levels of transcriptional activation with little evidence for H3K4me3 and H3K36me3 modification

As illustrated above, it has been commonly observed that the H3K4me3 modification at promoters and the H3K36me3 modification within transcription units reflect a past or present transcriptional activity of the corresponding genes. To probe whether this correlation is universal, we queried our data sets for genes that were transcriptionally active but displayed little H3K4me3 and H3K36me3 modification. We thus identified an unusual cluster of genes — the serum amyloid A (SAA) genes *Saa1, Saa2*, and *Saa3* — with such properties.

SAA proteins are apolipoproteins associated with high-density lipoprotein (HDL) in blood plasma. In mice, there are four principal liver-synthesized SAA proteins: SAA1, SAA2, SAA3, and SAA4, of which three, SAA1–SAA3, are synthesized during the acute phase of inflammation. In contrast, SAA4 is constitutively synthesized in the liver. The genes encoding these proteins are between 2.5 to 4.5 kb long and each has four exons.

The genomic response of these genes is shown in Fig. 7; Fig. 7a shows profiles of Pol II and histone H3 modification as in Fig. 4 with quantitations shown in Fig. 7b. Consistent with their known expression patterns, the acute inflammatory *Saa1* and *Saa2* genes exhibit a large increase in Pol II occupancy within 4 h of PH, an increase that was also observed in the sham samples. The Pol II density then decreases by the next time point at 10 h, to return in waves at 44 and 72 h. The *Saa3* gene also displays a burst of activity at 4 h, albeit much less robust than that of the *Saa1* and *Saa2* genes, and does not display the later waves of Pol II occupancy (Fig. 7b). In contrast, the constitutive *Saa4* gene displays a relatively constant density of Pol II.

As expected, the constitutive *Saa4* gene displays a clear presence of H3K4me3 and H3K36me3 modified histones. In contrast, the *Saa1* and *Saa2* genes display very little H3K4me3 and H3K36me3 modification at any of the three times that they are heavily occupied by Pol II. This deficiency is unlikely to be owing to the very high levels of Pol II occupancy, because very highly transcribed genes (e.g., the albumin encoding *Alb* gene shown in Additional File 5: Figure S5) can be extensively H3K4me3 and H3K36me3 modified. The deficiency also does not require high levels of transcription, because the *Saa3* gene has a much lower level of Pol II occupancy — even lower than that of the *Saa4* gene — and does not display H3K4me3 and H3K36me3 modification. Interestingly, the *Saa1, Saa2* and *Saa3* genes all display evident H3K36me2 modification at 60 h, indicating that (i) H3K36 modification per se is not likely to be impeded and (ii) H3K36me3 methylation is not compromised by a lack of H3K36me2 modification. These observations indicate that related genes — the *Saa* genes — can display high levels of Pol II activity without association with two of the principal Pol II-associated histone modifications: H3K4me3 and H3K36me3.

## Discussion

We have used mouse liver regeneration following 70% PH to study genome-wide cycles of gene expression, with particular attention to the re-entry of quiescent differentiated cells — hepatocytes — into the cell-division cycle. We examined gene expression by RNA-Seq of poly(A)-containing RNA, to probe a “phenotypic” outcome of active transcription, and by ChIP-Seq of Pol II, and H3K4me3 and H3K36me3 to probe states of genome response. The interpretation of the RNA-Seq analyses was more robust than that of the ChIP-Seq analyses probably owing to (i) a lower sequence complexity in the starting material (i.e., transcriptome vs. whole genome sequence), (ii) a greater amplitude in signal response (i.e., each gene is only present once per genome), and (iii) the need for epitope purification in the ChIP-Seq analyses. Furthermore, as less tissue was required for RNA-Seq, we could perform separate analyses of each of the three livers pooled for the ChIP-Seq analyses, making for more robust statistical analyses.

### Multiple cycles of gene expression following PH

Fig. 8 shows a summary of the PCA gene-expression “cycles” that we observed following sham and PH surgery. There were two discrete post-PH responses: (i) an early dynamic state from 1–20 h largely shared with the sham samples and (ii) a second dynamic state from 28–72 h specific to the PH samples and which returned to the resting state by one week. Although we have studied the regenerating liver as a whole and have thus not distinguished among (i) different cell types (e.g., hepatocyte and non-hepatocyte), nor (ii) cells in different states (e.g., proliferative vs nonproliferative), the results probably largely reflect proliferating hepatocytes, as hepatocytes make up the large majority of liver cells and as proliferation is the major *de novo* activity in the regenerating liver.

**Figure 8.**
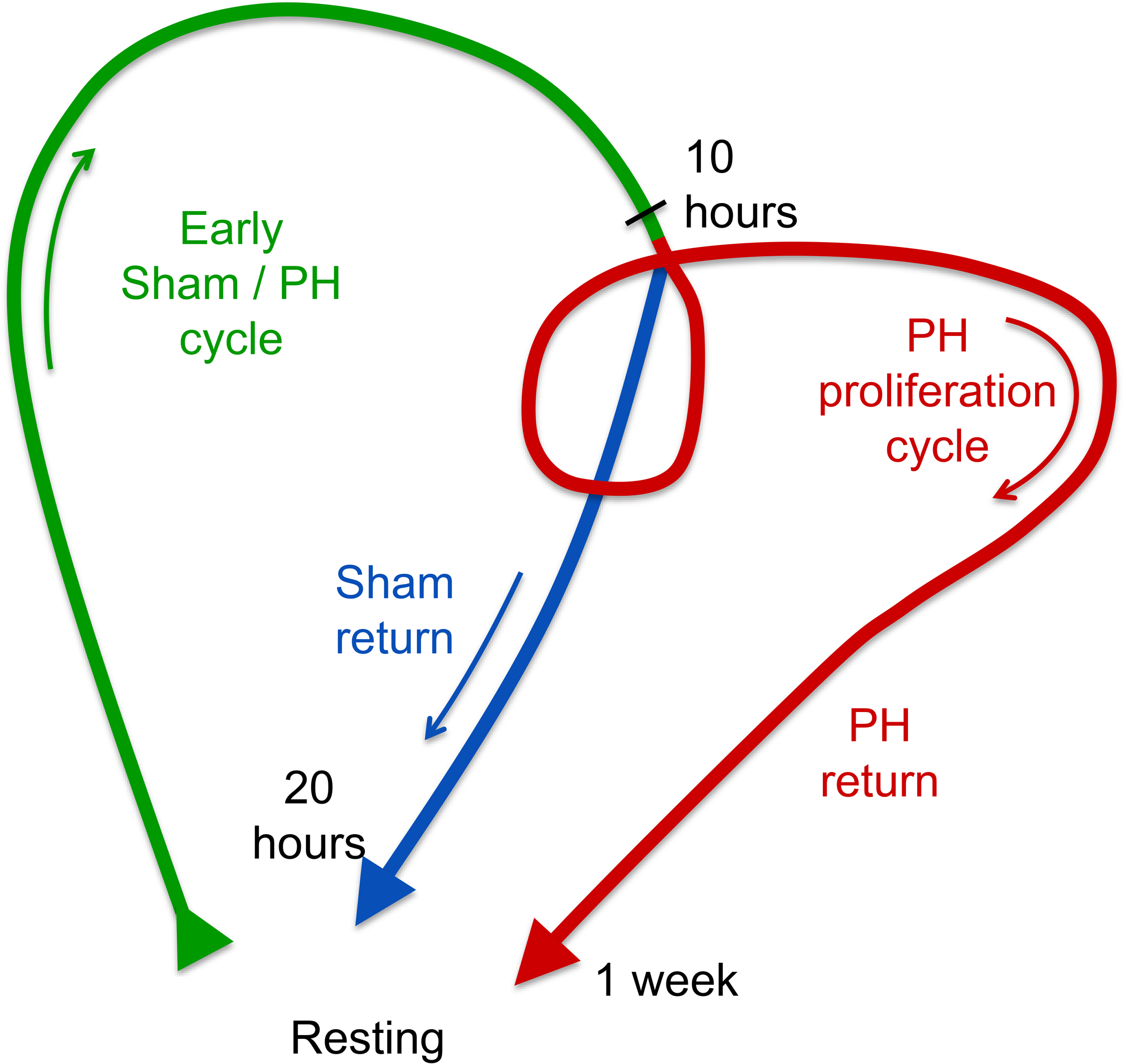
Graphic summary of cyclic gene-expression responses to sham and PH surgeries. Green, shared sham and PH surgery response; blue, sham-surgery-specific response; red, PH-surgery-specific response. See text for details.

The early, shared sham and PH cycle was surprisingly long before the sham program returned to a near resting state around 20 h and the PH program entered the PH-specific dynamic state. This second state probably reflects hepatocyte proliferation and as such multiple rounds of cell division. For example, examination of the RNA levels for the cell-cycle rich Set 3.7 (Fig. 2) or for the cell-division-cycle genes themselves (Fig. 3) reveals separate peaks of expression at 44–48 h and 72 h, most likely corresponding to the first and second rounds (or waves) of hepatocyte division.

We hypothesize that the gene-expression program in the sham samples is owing to a robust response to anesthesia, painkiller, skin wounding, and/or stress as a result of the surgical insult. A prominent feature of this sham gene-expression pattern is the degree to which cell-division-cycle genes are involved even though ultimate cell proliferation is absent (see Fig. 3). This ultimately nonproductive cell-proliferation response may be related to the selective advantage that liver regeneration has had during evolution. Indeed, liver regeneration probably evolved to allow organisms to survive the otherwise lethal effects of, for example, ingested toxins that can overwhelm and kill hepatocytes during the process of detoxification (see [6]). The full extent of eventual hepatocyte death is likely not evident upon intoxication. We hypothesize that given the ambiguity of the ultimate outcome, the liver evolved to prepare for the worst by immediately initiating an early program of cell-division-cycle gene expression, with a decision taken later on as to whether to progress through the cell-division restriction point [18] and onto S phase. It is also possible, however, and not mutually exclusive that G1-phase cell-division-cycle genes play specific roles beyond ones dedicated to cell proliferation. For example, Cyclin D1 is implicated in repression of hepatic gluconeogenesis [19], which may play a role in a damage response.

### What are the PH responses?

Among the 12,025 different transcripts we detected, one half — Set 2 — exhibited little change in abundance after PH. We suggest that the Set 2 transcripts largely represent RNAs that continue to be synthesized in both proliferating and non-proliferating cells. Consistent with this suggestion, Set 2 is enriched in transcripts encoded by genes involved in ubiquitous cellular processes such as macromolecular metabolism (e.g., RNA synthesis, regulation of gene expression; Additional File 2: Figure S2b). Among the transcripts that change in abundance during liver regeneration (i.e., Set 3), those that decrease generally represent genes involved in normal liver functions and may decrease because only the subset of non-proliferating hepatocytes maintain their expression. In contrast, those transcripts that increase in abundance, being largely involved in cell growth and proliferation, probably represent *de novo* functions that take place in only the subset of proliferating cells.

### Genomic responses to PH

To analyze genomic responses to transcriptional activation, we focused on Pol II occupancy and H3K4me3 and H3K36me3 modification at *de novo* activated genes following PH (so-called Post-PH genes); many Post-PH genes were cell-cycle genes. This analysis complements the study of [20], which examined the relationship of specific transcription-factor promoter (i.e., Cebpa and Cebpb) and Pol II gene occupancies during liver regeneration. In our study, for Post-PH genes, the presence of Pol II, and H3K4me3 and H3K36me3 modifications appeared in related waves. Importantly, Pol II was absent at 0 h and *de novo* recruited to promoters at 10-20 h well before gene-body transcription was fully active at 36–44 h. This pattern contrasts with Pol II occupancy in cells proliferating in culture, in which Pol II often occupies promoters constantly and an apparent key regulatory step is its release from the promoter region as an elongating Pol II [21, 22, 23]. In the instance of hepatocytes re-entering the cell-division cycle, however, *de novo* recruitment of Pol II to promoters is clearly temporally separate from Pol II promoter release. These results further emphasize the importance of paused-Pol II promoter release — and all its associated activities (e.g., Pol II phosphorylation) — as a regulatory step.

The two histone modifications analyzed here displayed very different patterns. As previously described (reviewed by [5]), the H3K36me3 modification was associated with active transcription. In contrast, H3K4me3 was evident at the TSS of Post-PH genes in the resting liver and increased only to a limited extent post PH as transcription was activated. These results contrast with those of [24], who specifically detected, at three days of liver regeneration, newly acquired H3K4me3 modifications at over 4000 sites of which only a minority were associated with increased gene transcripts; this difference may reflect the induction of liver regeneration by a non-PH protocol involving portal vein branch ligation. The role of the H3K4me3 mark in transcriptional regulation is not known [25]. The patterns of H3K4me3 modification observed at promoters of cell-proliferation genes in quiescent hepatocytes are consistent with the idea that H3K4me3 modification provides a memory of prior activity — that is from when the hepatocytes were proliferating during development to form the liver. Consistent with this hypothesis, we see a general absence of H3K4me3 at TSS of genes not related to liver function or development (e.g., neuronal genes like myelin, etc.) — genes that were likely never active in these cells or their progenitors.

### Comparison of genomic responses to two different gene expression cycles: circadian and cell division

We compared gene expression changes in response to the circadian [1] and cell-division (this study) cycles. These two cycles differ profoundly in their context. Whereas adult mouse hepatocytes undergo an essentially uninterrupted lifelong series of circadian cycles, these cells rarely enter the cell-division cycle. Thus, for genes regulated by the circadian cycle there is at most a 24 h break between times of activation, whereas for the cell-division cycle described here — for a highly differentiated tissue — many genes devoted to cell proliferation have been silent for weeks before activation by PH.

As previously noted [9] and illustrated here, while post-PH liver regeneration induces a dramatic shock to the hepatocyte gene-expression program, the circadian cycle continues unperturbed. The circadian cycle probably continues in all hepatocytes, whether proliferating or not, because the overall cycling levels of mRNAs encoding the master circadian regulators *Arntl* and *Nrd1d1* do not change during regeneration. Thus, here, the circadian cycle is resistant to major changes in cell gene-expression status.

To identify differences in genomic responses in the circadian and *de novo* cell-division cycles, we compared circadian and PH-specific Post-PH changes ([1]; this study). There were differences in the appearance of Pol II at promoters and gene bodies as well as H3K4me3- and H3K36me3-containing nucleosomes. For genes that are circadian-cycle regulated, Pol II levels at the promoter and gene body cycled in parallel indicating that release of a “paused” Pol II is not a regulatory step [1]. In contrast, the Post-PH genes in liver regeneration displayed promoter-bound Pol II significantly before Pol II could be detected in gene bodies. Perhaps, owing to the rhythmic and frequent activation and repression of gene transcription during the circadian cycle, there is a rapid transition from promoter-bound to elongating Pol II.

Modulation of H3K4me3 and H3K36me3 levels also differed between the circadian and cell-division cycles, for there was less variation, particularly for the H3K36me3 mark, during the circadian cycle than during the cell-division cycle. Such less variation could be owing to longer than 24-h periods being required for the disappearance of these methylation marks. Thus, it is the genes that have not been transcribed for a very long timespan that display the most conspicuous differences, in particular for the H3K36me3 mark.

### The H3K36me3 mark

In liver regeneration, the H3K36me3 mark displays a very different pattern from H3K4me3. H3K36me3 is absent from Post-PH genes until there is Pol II occupancy within the gene body at 36–72 h post PH; a pattern consistent with the association of the sole H3K36me3 methyltransferase SetD2 with the traveling Pol II [26, 27, 28, 29, 30, 31]. As aforementioned, H3K36me3 appears at the first internal exon. Clearly, H3K36me3 deposition requires H3K36me2, which is associated with active transcription units [17]. We noted that past the first internal exon genes with strong H3K36me3 signals possessed relatively little H3K36me2 modification, whereas at genes with less H3K36me3 signal the two modifications generally co-existed (Fig. 6). We imagine two explanations for how the H3K36me2 and H3K36me3 modifications might co-exist in the regenerating liver: The first explanation is that there is a mix of cells (e.g., different cell types or cells such as hepatocytes in different proliferative states) possessing genes with different H3K36me2 and H3K36me3 modification states. The second explanation is that individual genes have a mix of H3K36me2 and H3K36me3 modified nucleosomes, and that the level of H3K36me3 may be proportional to the amount of productive transcription of a given gene. Conceivably, for genes with mixed H3K36me3 signals, SetD2 has not yet saturated the H3K36me2 modification, perhaps owing to insufficient transcriptional activity.

### Unusual histone modification profiles

While examining our data sets for the general correlation between H3K4me3 or H3K36me3 methylation and transcription, we uncovered exceptions where transcribed genes had little of these two histone modifications. They were not many and they were often but not necessarily highly transcribed genes. The clearest example was the collection of related *Saa* genes shown in Fig. 7. Here, we found the highly transcribed *Saa1* and *Saa2* genes as well as the less highly transcribed *Saa3* gene to display little H3K4me3 or H3K36me3 modifications even though the H3K36me2 modification at 60 h was readily evident. These observations emphasize that the H3K4me3 and H3K36me3 histone modification marks are not ubiquitous in the transcription of all genes. That related genes — *Saa1, Saa2, and Saa3* — but not a third — *Saa4* — should display the same lack of histone modification is curious and suggests an evolutionarily conserved gene expression response.

## Conclusions

This study reveals a surprising early parallel response of the mouse liver to either PH or simple sham operatory procedures, indicating that anesthesia, painkiller, skin wounding, or stress are the first major effector of a liver gene-expression response to PH. Surprisingly, this early non-PH-specific response includes many cell-division-cycle genes even though there is no significant cell proliferation when no PH is performed. Thus, PH has an apparent delayed effect that results in full cell-division-cycle entry. In contrast to cell culture, where cell division is persistent, PH results in the proliferation of cells — adult hepatocytes — that have long been non-proliferative. Under such conditions of cell-division-cycle re-entry, transcriptionally activated genes are generally already modestly marked at the TSS by H3K4me3. This mark increases alongside Pol II TSS recruitment as the PH response proceeds, followed by Pol II transcription elongation and H3K36me3 appearance on a foundation of H3K36me2 modification after the first internal gene exon when transcriptionally fully active. Furthermore, we identified unusual histone mark patterns, whereby highly transcribed genes were devoid of H3K4me3 or H3K36me3 marks.

## Methods

Extended experimental methods are described in supplemental material (Additional File 10: Supplementary Material).

### PH and sham surgical procedures

Mouse entrainment and surgical procedures were as described [10]. Briefly, 12–14 week-old C57/BL6 male mice were used for PH or sham surgery after four weeks entrainment: two weeks with a ZT0–ZT12 light and ZT12–ZT24 dark circadian cycle followed by two weeks of ZT0–ZT12 light with fasting and ZT12–ZT24 dark with feeding. Three mice at a time were subjected to 2/3 PH at ZT2 as described [32] and sacrificed together at either 1, 4, 10, 20, 28, 36, 44, 48, 60, or 72 h or one or four weeks post-surgery. Sham-operated controls — three at a time — were subjected to laparotomy and sacrificed at 1, 4, 10, 20 or 48 h post surgery.

### RNA-Seq library preparation and sequencing

Selected poly(A)-containing mRNA from individual livers was used to prepare strand-specific libraries for 100 nucleotide single-end sequencing with an Illumina HiSeq 2100 machine.

### ChIP, ChIP-Seq library preparation, and ultra-high-throughput sequencing

The three livers per time point were pooled for analysis. The ChIP protocol was adapted from [1] (see Additional File 10: Supplemental Material). For each ChIP, 19 μg sonicated mouse liver DNA was mixed with 1 μg sonicated human HeLa cell DNA for ‘spiking’ [33]. 10 ng ChIP DNA was used for 100 nucleotide Illumina HiSeq 2100 paired-end sequencing after library preparation with 14 cycles of PCR amplification and no electrophoretic size selection.

### RNA-Seq and ChIP-Seq data preparation and quantification

RNA-Seq and ChIP-Seq results were analyzed as described in the Supplemental Methods (Additional File 10: Supplementary Material). For paired-end ChIP-Seq, the terminal 50 bp sequences were mapped onto the mouse (mm9) and human (hg19) genomes with Elandv2e and the Ensembl 67/NCBI 37 transcription-unit annotation was used. Among the multiple transcription units associated with a given gene, only the one containing the maximum promoter-associated Pol II occupancy was used for the analysis of Pol II body, H3K4me3 and H3K36me3 signals.

### Transcriptome analyses

Transcripts with significant accumulation (12,025) were classified into stable and changing expression. Only transcripts with associated corrected p-values lower than 1 × 10^−7^, and log2 fold-changes higher than 0.5 or lower than −0.5 were retained. Dendogram, PCA, and Set 3 PAM clustering analyses, gene expression profiles, post-PH- and sham-sample comparisons, functional enrichment and pathway annotation are described in the Supplemental Methods (Additional File 10: Supplemental Material).

### H3K36me2 and H3K36me3 density profiles at a one-nucleotide resolution

Nucleotide positions 4 kb upstream and 1 kb downstream of the 3′ end of the first internal exon were analyzed. Single nucleotide densities using the central 50 bp of sequenced fragments were quantified. A matrix of quantifications per Transcription Unit (TU) was built and analyzed on the R statistical software [34]. Z-scores were calculated from the quantifications and displayed with the heatmap.2 function included on the ‘ggplots 2.14.1’ [35].

## Abbreviations

C: Control sample
GO: Gene Ontology database
H3: Histone 3
H3K4me3: trimethylated histone H3 lysine 4
H3K36me3: trimethylated histone H3 lysine 36
H3K36me2: dimethylated histone H3 lysine 36
HDL: High-density lipoprotein; mRNA: Messenger RNA
PAM: Partitioning Around Medoids clustering
PC: Principal component
PCA: Principal component analysis
PH: Partial hepatectomy
Pol II: RNA polymerase II
RNA-Seq: ultra-high-throughput RNA-sequencing
RNA-Seq: ultra-high-throughput DNA-sequencing
RPKM: Reads per Kb of transcript per million mapped reads
S: Sham surgeries
Saa: Serum amyloid A
TSS: Transcriptional start site
TU: Transcription Unit
X: Excised liver hepatectomy
ZT: Zeitgeber Time

## Declarations

### Ethics approval and consent to participate

Not applicable

### Consent for publication

Not applicable

### Availability of data and material

All the datasets generated in this study are available in the NCBI Gene Expression Omnibus (GEO; http://www.ncbi.nlm.nih.gov/geo) under the accession number GSE95136.

### Authors’ contributions

The experiments were conceived and designed by LR, DV, NH, NG, and WH. The experiments were performed by DV. LR, VP, NG and WH analyzed the data. LR, NG, and WH wrote the manuscript. All authors participated in the discussion of the data and in production of the final version of the manuscript.

## Acknowledgements

We thank S. Offner for advice on the partial hepatectomy operation; the Lausanne Genomic Technologies Facility for the RNA-Seq library preparation and high-throughput sequencing, L. Fajas for a critical reading of the manuscript. We thank Keith Harshman, Director of the Lausanne Technologies Facility, where all the ultra-high-throughput sequencing was performed, and Ioannis Xenarios, Director of the Vital-IT (http://www.vital-it.ch) Center for High Performance Computing of the Swiss Institute of Bioinformatics.

We thank the CycliX Consortium: Nouria Hernandez,^1^ Mauro Delorenzi,^2,3,4^ Bart Deplancke,^5^ Béatrice Desvergne,^1^ Nicolas Guex,^6^ Winship Herr,^1^ Felix Naef,^5^ Jacques Rougemont,^7^ Ueli Schibler,^8^ Teemu Andersin,^8^ Pascal Cousin,^1^ Federica Gilardi,^1^ Pascal Gos,^8^ Fabienne Lammers,^1^ Maykel Lopes,^1^ François Mange,^1^ Shilpi Minocha,^1^ Sunil Raghav,^5^ Dominic Villeneuve,^1^ Roberto Fabbretti,^6^ Volker Vlegel,^6^ Ioannis Xenarios,^1,2,6^ Eugenia Migliavacca,^1,6^ Viviane Praz,^1,2^ Fabrice David,^2,7^ Yohan Jarosz,^2,7^ Dmitry Kuznetsov,^6^ Robin Liechti,^6^ Olivier Martin,^6^ Julien Delafontaine,^2,7^ Julia Cajan,^5^ Cristian Carmeli,^3^ Kyle Gustafson,^1^ Irina Krier,^5^ Marion Leleu,^2,7^ Nacho Molina,^5^ Aurélien Naldi,^7^ Leonor Rib,^1^ Jonathan Sobel,^5^ Laura Symul,^5^ Gergana Bounova1,^2^ and *Philippe Jacquet^2,7^

^1^Center for Integrative Genomics, Faculty of Biology and Medicine, University of Lausanne, 1015 Lausanne, Switzerland

^2^Swiss Institute of Bioinformatics, 1015 Lausanne, Switzerland

^3^Bioinformatics Core Facility, Swiss Institute of Bioinformatics, 1015 Lausanne, Switzerland

^4^Department of Oncology and Ludwig Center for Cancer Research, Faculty of Biology and Medicine, University of Lausanne, 1011 Lausanne, Switzerland

^5^Interfaculty Institute of Bioengineering, School of Life Sciences, Ecole polytechnique Fédérale de Lausanne, 1015 Lausanne, Switzerland

^6^Vital IT, Swiss Institute of Bioinformatics, 1015 Lausanne, Switzerland

^7^Bioinformatics and Biostatistics Core Facility, School of Life Sciences, Ecole polytechnique Fédérale de Lausanne, 1015 Lausanne, Switzerland

^8^Department of Molecular Biology, Faculty of Sciences, University of Geneva, 1211 Geneva, Switzerland

## Additional Files

### Additional File 1

**Title**: Supplemental Figure S1

**File format**:

pdf

**Description of the data**:

(**A**) Description of the mouse liver regeneration stages, collection time points of liver samples and the food and light conditions of the mice. Adapted from [10]. X = eXcised liver hepatectomy; C = controls; S = sham surgeries; W = week

(**B**) Number of replicates per experimental condition and the associated RNA-Seq and ChIP-Seq experiments performed. In a preliminary PH analysis (Series 1), we performed ChIP-Seq on pools of three livers at the post-PH time points indicated and no sham surgeries. Subsequently, we performed Series 2 with the full set of time points and selected sham surgeries, with RNA-Seq analyses of generally three individual livers and, where indicated, ChIP-Seq on pools of three livers. Note that Series 2 was performed in three separate time periods (Series 2.1, 2.2, 2.3); for two samples, X20 and X36, duplicate samples were prepared in Series 2.1 and 2.2. The samples from Series 2.1 were used in the data shown and discussed in the text. The Series 1 60 h sample was used for the K36me2 vs. H3K36me3 study in Fig. 6.

(**C**) Correlation of three triplicates from each of Series 2.1, 2.2, and 2.3 (nine samples total) at time C0. The replicates are indicated on the diagonal. Above the diagonal, pairwise scatterplots display the similarity between replicates, and below the diagonal, each pairwise Pearson coefficient is indicated. Correlations within experimental series (Pearson correlation coefficients of at least 0.98) show slightly better correlation coefficients than among series (Pearson correlation coefficient of at least 0.94).

(**D**) Two-dimensional plot displaying the coordinates of the collected samples in the PC1 and PC2 of the PCA using the set of 12,025 expressed genes as shown in Fig. 1b, but with the standard deviations in PC1 and PC2 of the replicates for each condition displayed as ovals.

### Additional File 2

**Title**: Supplemental Figure S2

**File format**:

pdf

**Description of the data**:

(**A**) Distribution of silhouette scores as a result of the PAM clustering of the varying gene expression Set 3 into 2 to 13 groups (k). The clustering into 7 groups was retained for our analyses in the main text (Fig. 2). In this Set3.1 to 3.7 clustering, set 3.7 contained a large proportion of cell-cycle genes. The set in the k = 2–6 and 8–12 PAM clusterings most like Set 3.7 is labeled “3.7-like” in each case. For each clustering, the number of genes (nj) per group (j) is indicated to the right together with the average silhouette score (ave_iϵCj_ Si). To the left of each clustering, the number of Set 3.7 genes from k = 7 in each group is indicated. Across clusterings, the highest average silhouette score is found for the most 3.7-like sets of genes.

(**B**) Summary of the results of the functional enrichment analysis on the RNA-seq Set 1, Set 2 and Set 3 results. The GO terms displaying an enrichment p-value lower than 10E-10 were kept for analysis with the REVIGO tool. REVIGO aggregates synonymous GO terms and displays the aggregated terms as circles where the distance among circles indicates their similarity within the GO structure and their color indicates the associated p-value, with blue signifying the lowest p-values. Selected GO terms with highest p-values are shown with the circle aggregates. Below, the highlighted GO terms have been listed with their associated p-values with a log10 scale.

(**C**) Gene expression patterns post PH in the KEGG cell-cycle pathway. The gene nodes in the KEGG cell-cycle pathway were colored using the ‘pathview’ R package. Set 1 genes are colored gray, Set 2 genes are colored yellow and Set 3 genes are displayed as a heat map that shows the relative transcript abundance between 0 h and 4 weeks post PH from Fig. 2a. For the twenty-seven KEGG cell-cycle pathway nodes shared by multiple genes only the pattern for a representative gene (identified in Additional File 2: Figure S2d).

(**D**) Selection of the representative gene within shared KEGG cell-cycle nodes. The names of the shared KEGG cell-cycle nodes are indicated with the node-associated genes indicated with their respective RNA-Seq set. Within nodes, genes with differential expression post PH were favored. Otherwise genes with the highest gene expression within a node were selected. The final selections are highlighted in green.

### Additional File 3

**Title**: Supplemental Figure S3

**File format**:

pdf

**Description of the data**:

(**A–E**) Additional examples of transcriptional gene activity post PH. Displays of the ChIP-Seq results of five different gene sets — (A) *Bcl2l1* and *Tpx2*; (B) *Cdca2* and *Kctd9*; (C) *mKi67*; (D) *Ppp1r12b* and *Ube2t*; and (E) *Rad51ap1* and *D6Wsu163e* — are shown as described in Fig. 3.

### Additional File 4

**Title**: Supplemental Figure S4. Pol II, H3K36me2 and H3K36me3 densities at 60 h post PH for the *Cebpa* gene.

**File format**:

pdf

**Description of the data**:

(**A**) Genomic view of the *Cebpa* gene. Densities of the central 50 bp of paired-end reads for Pol II (pink), H3K36me3 (dark blue) and H3K36me2 (light blue) ChIP fragments are shown for 60 h post PH. Similarly, input fragments at 60 h post PH are shown.

### Additional File 5

**Title**: Supplemental Figure S5. Pol II, H3K36me2 and H3K36me3 densities at 60 h post PH for the *Alb* gene.

**File format**:

pdf

**Description of the data**:

(**A**) Genomic view of the *Alb* gene. Densities of the central 50 bp of paired-end reads for Pol II (pink), H3K4me3 (green) and H3K36me3 (blue) ChIP fragments are shown for 0 h to 1 week post PH. Similarly, densities for H3K36me2 and input fragments at 60 h post PH are shown.

(**B**) H3K4me3 (green), Pol II promoter (burgundy) and body (pink), and H3K36me3 (blue) log2 ChIP/Input fragment-density comparison for the *Alb* gene post PH. The central 50 bp of the paired-end reads were used for quantification. The regions used for each quantitation are given in the text.

### Additional File 6

**Title**: Supplemental Table S1

**File format**:

xlsx

**Description of the data**:

List of genes in Set 2

### Additional File 7

**Title**: Supplemental Table S2

**File format**:

xlsx

**Description of the data**:

List of genes in Set 3 with their associated classification in Sets 3.1 to 3.7 plus the averaged RPKMs per time point in log2 scale.

### Additional File 8

**Title**: Supplemental Table S3

**File format**:

xlsx

**Description of the data**:

Complete list of GO terms associated to genes in Sets 1 to 3.7 with the associated corrected p-value.

### Additional File 9

**Title**: Supplemental Movie S1

**File format**:

mpeg

**Description of the data**:

Temporal changes in gene expression post PH in the KEGG cell-cycle pathway. Animated expression changes post PH in the KEGG cell-cycle pathway. At each time point between 0 h and 4 weeks the gene nodes in the KEGG cell-cycle pathway were colored using the ‘pathview’ R package [13]. Genes are colored as in Additional File 2: Figure S2d.

### Additional File 10

**Title**: Supplemental Material

**File format**:

pdf

**Description of the data**:

Supplemental Methods and References

